# NFAT5 Confers Differential Expression in Innate Immune Cells vs. Renal Epithelial Cells in the Face of Osmotic Stress

**DOI:** 10.1101/2025.11.18.689108

**Authors:** Xander Kraus, Favour Nnah, Craig Cowley, Patricio Araos, Brandi M. Wynne

## Abstract

The kidney develops an osmotic gradient, up to 1300 mOsm, to drive the reabsorption of water. However, this creates a harsh environment for renal epithelial and immune cells. These cells need to develop a survival mechanism to prevent against osmotic stress and maintain homeostasis in this harsh environment. <u>N</u>uclear <u>F</u>actor of <u>A</u>ctivated <u>T</u> Cells 5 (NFAT5, also known as <u>Ton</u>icity-Responsive <u>E</u>nhancer-<u>B</u>inding <u>P</u>rotein factor or TonEBP) is a master transcription factor that responds to changes in osmolarity, regulating the transcription of genes that control the intracellular accumulation of osmolytes. Studies suggest that NFAT5 may be responsible for cell survival in this environment. Both epithelial and immune cells play a role in pro-inflammatory responses and disease progression. Additionally, NFAT5 may also play divergent roles, promoting the expression of pro-inflammatory genes in immune cells, and anti-inflammatory genes in epithelial cells. We hypothesized that NFAT5 expression was protective in epithelial cells, while stimulation in immune cells may be proinflammatory. We performed survival assays to assess an NFAT5-dependent role in hyperosmolar environments. We also assessed gene expression and identified targets that may be responsible for cell survival, as well as demonstrating divergent mechanisms in epithelial and immune cells. In conclusion, we found NFAT5 is critical to cell survival in hyperosmolar environments and identified transcripts such as *bgt1, aqp1 and akr1b3* that may explain the difference in survival. We also observed a difference in expression between epithelial and raw cells, including the same targets thought to protect against hyperosmotic stress in IMCD3 cells that showed differential regulation by NFAT5. These data suggest a divergent NFAT5 mechanism in two cell types present in the kidney.

## Introduction

The kidney functions to maintain water and electrolyte balance, which includes maintaining blood pressure. To do this, the kidney develops a concentration gradient from the isosmotic cortex through the outer medulla (OM) to the inner medulla (IM). Developing this gradient is mandatory for water reabsorption and concentrating urine. This is primarily established by the selective reabsorption of sodium in the IM along with the counter-current mechanism allowing for urea recycling in the medullary Loop of Henle. Together, these mechanisms induce a force for collecting duct (CD)- dependent reabsorption of water and allows for IM osmolyte concentrations of up to 1300 mOsms in humans and 4000 mOsms in rodents^1^. Although this process is critical for concentrating urine, it poses a threat to cells, including renal epithelial cells and resident immune cells, along this gradient^2,3^. As such, these cells need a mechanism to survive in the hostile hyperosmolar environment, which includes increasing intracellular osmolytes such as betaine, taurine, sorbitol and myo-inositol which can help to prevent osmotic stress ^1,4^.

Previous studies have established <u>N</u>uclear <u>F</u>actor of <u>A</u>ctivated <u>T</u> Cells 5 (NFAT5, also known as <u>Ton</u>icity-Responsive <u>E</u>nhancer-<u>B</u>inding <u>P</u>rotein factor TonEBP) as a master transcription factor critical in the programmed response to hyperosmolar stress, specifically in epithelial cells^3^. With increasing extracellular osmolarity, NFAT5 induces expression of genes important for osmolyte and water transport. The importance of this gene is underscored by global deletion studies that demonstrate renal atrophy and increased mortality in that model^5,6^. Additionally, other studies have suggested that NFAT5 can also regulate non-hyperosmolar induced gene expression^3,4^. Studies have also shown that NFAT5 knockout (KO) both *in vitro* and *in vivo* leads to increased acute kidney injury markers and pro-inflammatory markers^1^. This is unsurprising, as NFAT5 belongs to the NFAT family of Rel-A homologues similar to other NFAT-associated transcription factors which primarily regulate immune cell function and cytokine signaling^7^. However, unlike other NFAT family members (NFAT1-4), NFAT5 uniquely responds to changes in osmolarity^7^. Single nucleotide polymorphisms (SNPs) in NFAT5 introns are associated with increased risk of inflammatory-related diseases, such as type 2 diabetes in humans^8–12^, and NFAT5 heterozygous mice have immune-related diseases^3,13,14^ suggesting a strong link between NFAT5 and inflammation. Yet, renal tubular-specific reduction in NFAT5 was shown to induce salt-sensitive hypertension^15^. It seems that whether this association exacerbates or alleviates the pathophysiology of the disease process fully depends upon the context of that specific disease.

Recent evidence indicates that NFAT5 exhibits fundamentally different functions depending on cell type and microenvironmental context^16,17^. In renal epithelial cells, NFAT5 functions primarily as an osmo-protective transcription factor, driving expression of osmolyte transporters and heat shock proteins that enable cell survival in the hypertonic renal medulla^18^. In contrast, in macrophages and other immune cells, NFAT5 promotes pro-inflammatory cytokine production and drives M1 macrophage polarization in response to both osmotic and non-osmotic stimuli^16,19^. This context-dependent activity appears to stem from differential reactive oxygen species sources and distinct co-factor interactions that direct NFAT5 to different gene promoters^20,21^.

The kidney is highly heterogeneous, including a vast network of innate immune cells^22^, along with renal epithelial cells- all of which responds to increasing interstitial osmolarity. For this study, we hypothesized that innate immune cells exhibit a profoundly different gene expression profile than IM renal epithelial cells and used two *in vitro* cell culture models, allowing us to look at differences in expression across cell lines. We hypothesize that NFAT5 is ultimately protective in renal epithelial cells via upregulation of key transporters yet promotes pro-inflammatory cytokines in immune cells.

## Methods

All drugs were purchased from Sigma Aldrich (St. Louis, MO), unless otherwise specified.

### Cell Culture

The inner medullary collecting duct cells (IMCD3) have been previously described and were a generous gift from Dr. Nirupama Ramkumar^23,24^. IMCD3 cells were grown in DMEM/F12 medium containing heat-inactivated fetal bovine serum (FBS, 10%) and penicillin/streptomycin (P/S, 1%) and plated on cell culture treated dishes. The RAW264 cell line (ATCC, Manassas VA) is a stable cell line used to study innate immune cells, such as macrophages, in cell culture. RAW cells were cultured in RPMI + GlutaMAX (ThermoFisher, Waltham MA) supplemented with non-heat inactivated FBS (10%) and pen/strep (1%). RAW cell lines are maintained on non-adherent cell culture dishes (non-treated) until plated for final experimental use, where standard cell culture treated dishes were used. RAW cells were passed using a gentle protocol of 0.48 mM EDTA in PBS until lifting (approximately 1-3 minutes) to preserve CD antigens. Both cell lines were grown at standard cell culture conditions (5% CO2/37^0^C).

### CRISPR Knockout

We generated a genomic NFAT5 KO IMCD3 line using a mixture of three short guide RNAs targeted at exon 5. As published previously^25^, per manufacturer’s instructions (Synthego, Menlo Park CA), IMCD3 cells were nucleofected with Cas9 ribonuclear protein. Single cell clones were developed from the nucleofected cells using limited dilution techniques. Western blot of NFAT5 demonstrated complete absence of NFAT5 (Fig 1).

**Figure 1.**
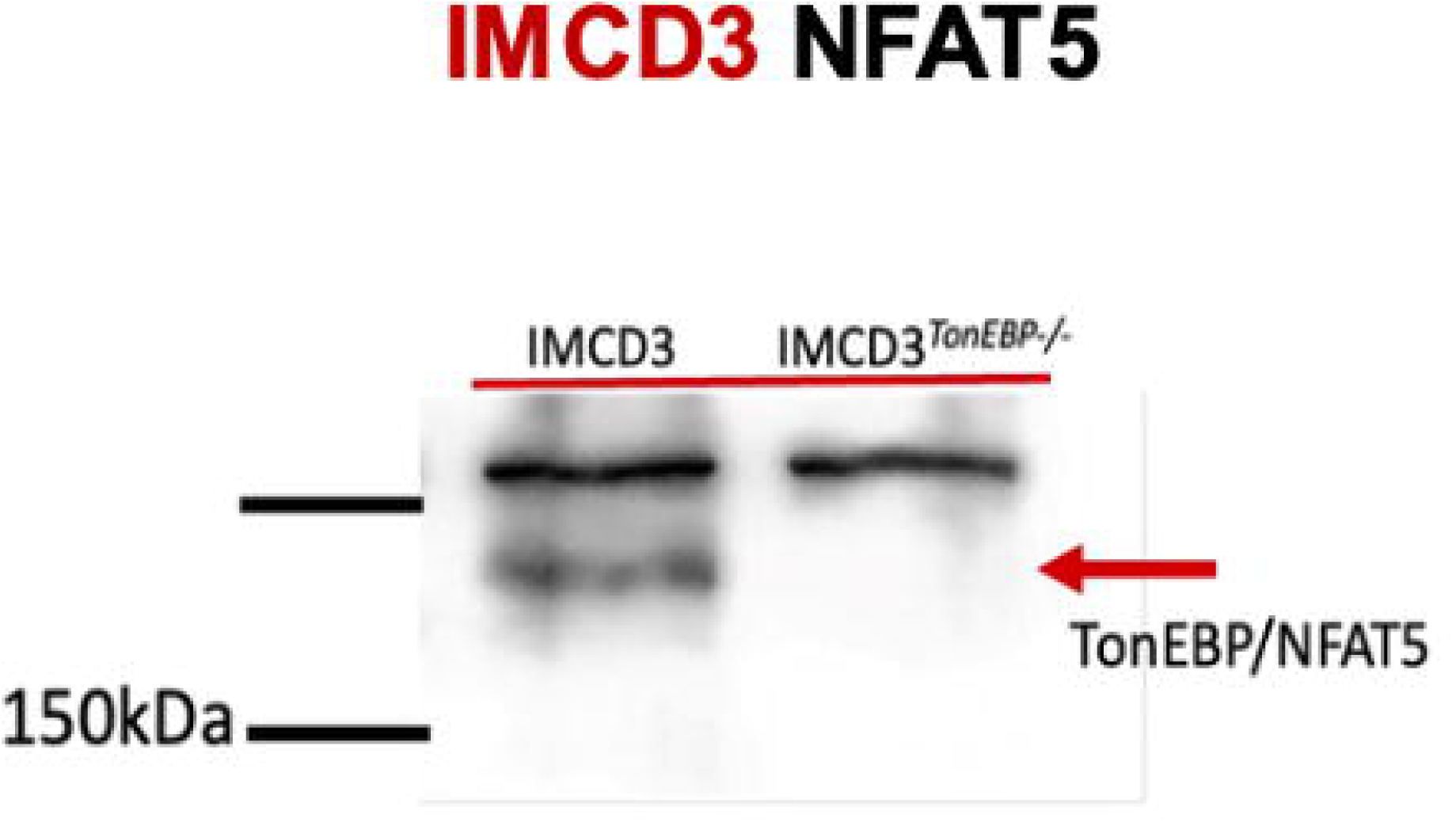
CRISP knockout of nuclear factor of activated t-cells (NFAT5) in inner medullary collecting duct (IMCD3) cells. NFAT5 (aka TonEBP) knockout was performed using a mixture of 3 short guide RNAs targeting exon 5 that were nucleofected with the Cas9 ribonuclear protein into murine inner medullary collecting duct (IMCD3) cells. Cell clones were collected and expanded. Total protein used for western blot showing the total NFAT5 knockdown.

### siRNA NFAT5 Knockdown

RAW 264.7 cells were transfected with siRNA against NFAT5 (or scramble, Lipofectamine RNAiMAX, ThermoFisher) in opti-MEM (Gibco).

### Quantitative PCR

Cells were plated in 6-well plates and grown with media containing additional osmolytes of additional sodium [+40mM/80mOsms or +80mM/160mOsms] or additional mannitol [+160mM/160mOsms] for (48hrs). Total mRNA was extracted (QIAGEN RNeasy Kit, Germantown MD) and used for real-time PCR (Luna Universal qPCR Master Mix) following cDNA library (LunaScript RT Supermix Kit, Alpharetta GA).

### Urea Stimulation

IMCD Wt (wild type) and NFAT5 knockout cells were grown in high urea to a maximum of 80 mM. Further increases were found to be non-viable. This concentration was established using step-wise increases in urea following stabilization for 24hrs at each concentration increase. Total mRNA was used for real-time PCR or to establish a cell survival curve.

### Cell survival curve

IMCD Wt and NFAT5 knockout cells were plated in 6-well plates. For cells not primed with urea, cells were grown for 48hrs before treating with vehicle (water) in media [approximately 320mOsm/kg], NaCl [+40mM/80mOsms or +80mM/160mOsms] or mannitol [+160mM/160mOsm]. After 0, 1, 3, and 5 days, cells were fixed (2% paraformaldehyde in PBS), permeabilized (0.5% Triton X-100 in PBS; 5 min) and incubated with Janus Green B dye (Molecular Probes, 0.2% in 1xPBS; 5 min).

IMCD Wt and NFAT5 knockout cells were plated in 6-well plates and allowed to grow for 24hrs. Cells were then primed with urea [80mM]. After another 24 hours, cells were further stimulated with either vehicle in media [approximately 320mOsm/kg] or NaCl [+40mM/80mOsms, +80mM/160mOsms or +120mM/120mOsms]. Cells were fixed (2% paraformaldehyde in PBS), permeabilized (0.5% Triton X-100 in PBS) before incubating with Janus Green B dye (Molecular Probes, 0.2% in 1xPBS; 5 min). Cell concentrations were assessed fluorometrically via plate reader (620nm). Cell concentrations from Wt vs. KO cells and treatments were plotted to show relative levels at all time points. The baseline growth curves of both Wt and NFAT KO cells (control media) were shown in all graphs, for representative demonstration of cell survival as compared to each experimental group; these were the same data.

### Statistical Analysis

Data were analyzed using Prism GraphPad 7 (San Diego, CA), using appropriate parametric or non-parametric analyses. All data are shown at mean ± SEM. Cell survival curves were analyzed by plotting a simple linear regression line; slopes were then compared. Different days were analyzed by two-way ANOVA, followed by post-hoc analysis. The fold change of mRNA expression between the untreated and ‘urea long stim’ was compared between the IMCD3 Wt and NFAT5 knockout using a *Students’* t-test. Significance between expression of treated and untreated IMCD Wt and NFAT5 knockout cells was tested with a 2-way ANOVA, followed by post-hoc analysis. Differences in the RAW 264.7 control and NFAT5 siRNA mRNA expression was compared using a *Students’* t-test.

### Primers

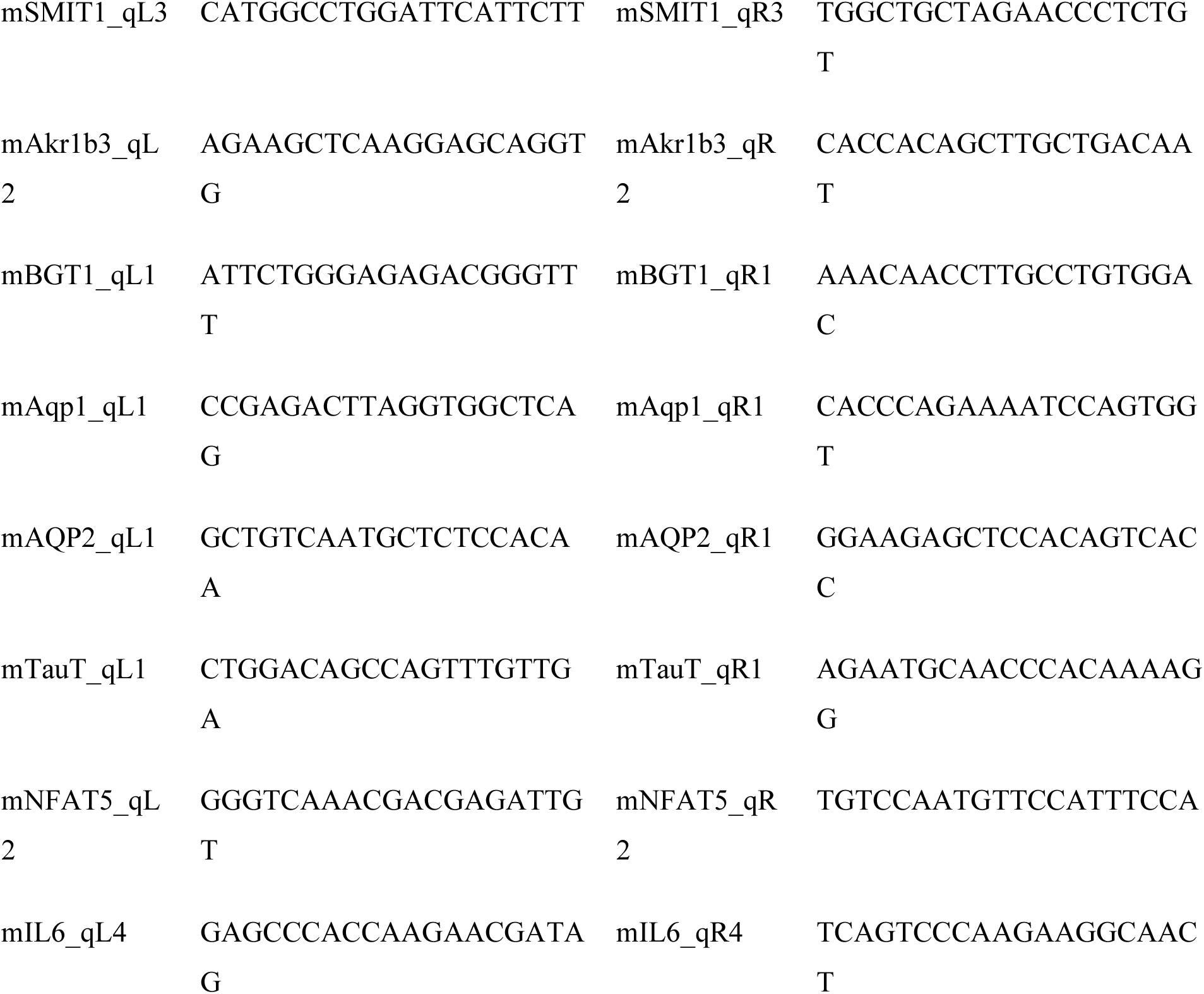
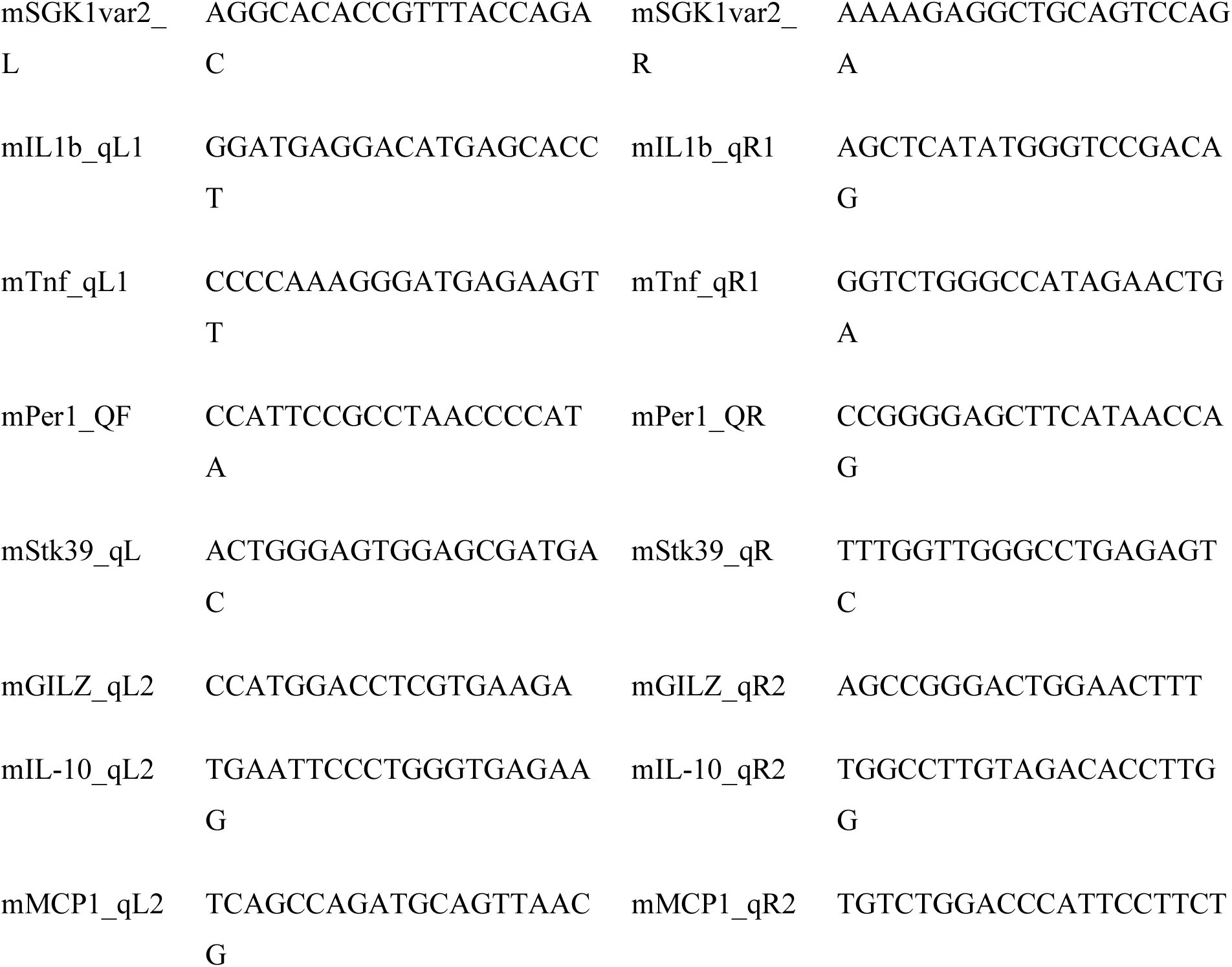

## Results

### Renal Epithelial Cell Survival Depends on NFAT5

Previous studies have demonstrated that both immune and epithelial cells exist in high osmolarity environments within the kidney^2^. Cells in this hostile environment need a mechanism to address the stress that comes with hyperosmolarity; however, no studies have specifically knocked down NFAT5 in these cell types *in vitro* and attempted to compare responses. NFAT5 is a transcription factor that regulates the cellular response to hypertonicity and seems to also play a role in immune cell function^3^.

Using a CRISPR KO method in IMCD3 cells (Fig 1), we assessed NFAT5-dependent long-term survival at increasing osmolarities over time, allowing the cells to acclimate to changes in osmolarity^25,26^. IMCD3 Wt cells and NFAT KO cells were treated with increased concentrations of NaCl [+40-80mM] or mannitol [+160mM] as an osmolar control, then fixed and stained with Janus Green B (Fig 2) to assess cell proliferation and survival. NFAT5 KO cells showed no reductions in cell survival in media alone [320mOsm/kg] compared to Wt counterparts. When presented with an additional [+40mM] NaCl (Fig 2A),

**Figure 2:**
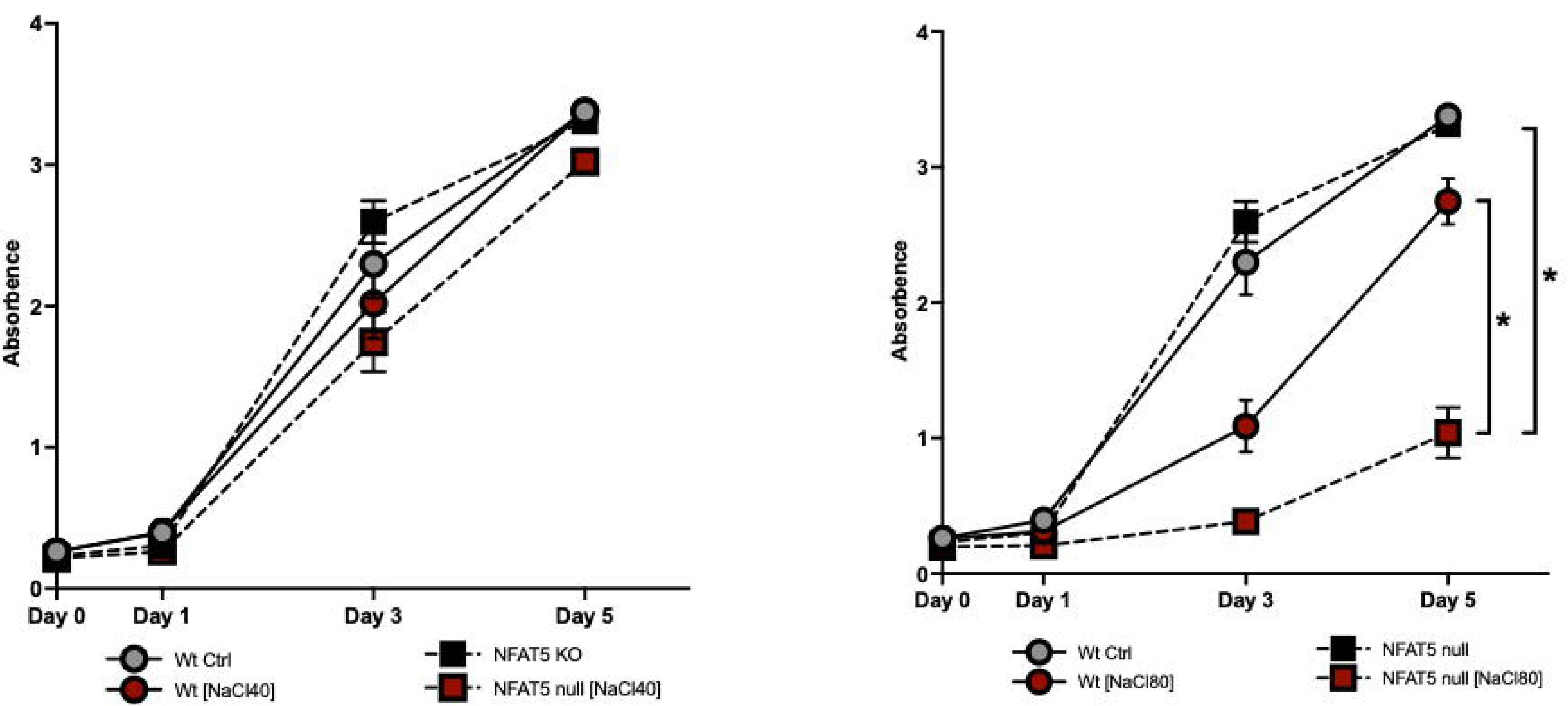
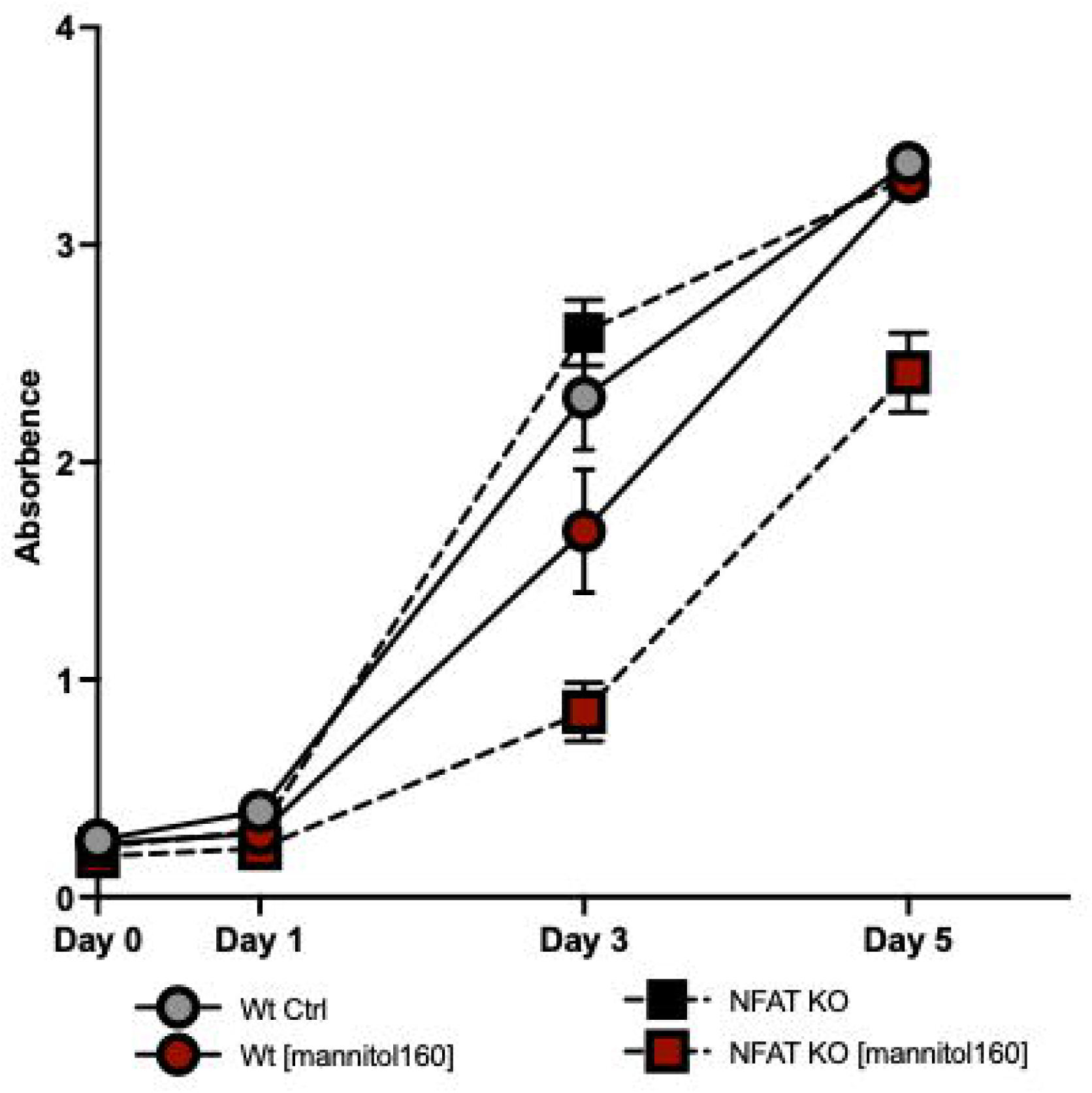
Nuclear factor of activated t-cells (NFAT5) is necessary for cell survival in hyperosmolar conditions. Cells were plated in media and then treated with vehicle or additional osmoles of: (A) NaCl [+40mM/80mOsm], (B) NaCl [+80mM/160mOsm] or (C) mannitol [+160mM/160mOsm]. Cell survival was determined at 0, 1, 3, or 5 days by staining with Janus Green B dye and measured at 620nm. spectrophotometer to determine concentration. Data were analyzed using a simple linear regression curve fitted to survival data followed by slope comparison, and are expressed as mean±SEM, n=4-5. *p<0.05, **p<0.01

NFAT5 KO cells exhibited no reduction in cell survival. With increasing NaCl [+80mM], Wt cells had a 47.4% reduction at day 3, that was not significant, (n=3-5). However, the NFAT5 KO cells exhibited a reduced cell survival at day 3 (16.5%) that recovered some, but overall survival significantly reduced (*p<0.05, n=4, linear regression slope) compared to both Wt cells in hyperosmolar NaCl [+80 mM] and NFAT5 KO cells in control media. Wt cells grown in the lower NaCl media [+40mM] (Fig 2B) or mannitol (Fig 2C) had no reductions in growth, and fully recovered by day 5 (NaCl 100.23%, mannitol 97.39% vs. control media, n=3-5), which was not seen in Wt cells in hyperosmolar NaCl [+80mM] (NaCl 81.33% vs. control media, n=5).

Given that NFAT5 KO cells in lower [40mM] concentrations of NaCl exhibited some recovery, we used a chronic ‘adaptive’ model, allowing the cells to recover before introducing an acute osmotic stimulus (Fig 3). Wt and NFAT5 KO cells were grown in media with urea increasing gradually over time, allowing the cells to gradually acclimate to hyperosmolarity. NFAT5 KO cells exhibited no differences compared to their Wt counterparts (Fig 3A). When given an additional [+80mM] NaCl neither Wt nor NFAT5 KO cells exhibited a significant reduction in cell growth, but there was a trending reduction (Fig 3B). Cells grown in urea with the highest NaCl [+120mM] fared worse, with NFAT5 KO cells incapable of any growth and significantly reduced compared to Wt. Surprisingly, Wt cell survival was not significantly reduced (Fig 3C). Interestingly, all cells exhibited a greater general survival ability with urea acclimatization. In a dose-dependent manner, IMCD3 NFAT5 KO cells could not recover from osmotic stress, thus confirming the importance for NFAT5 in osmoregulation. Yet, adaptation to urea suggests that there is an alternate method of responding to osmotic stimulus that may not be NFAT5 dependent.

**Figure 3:**
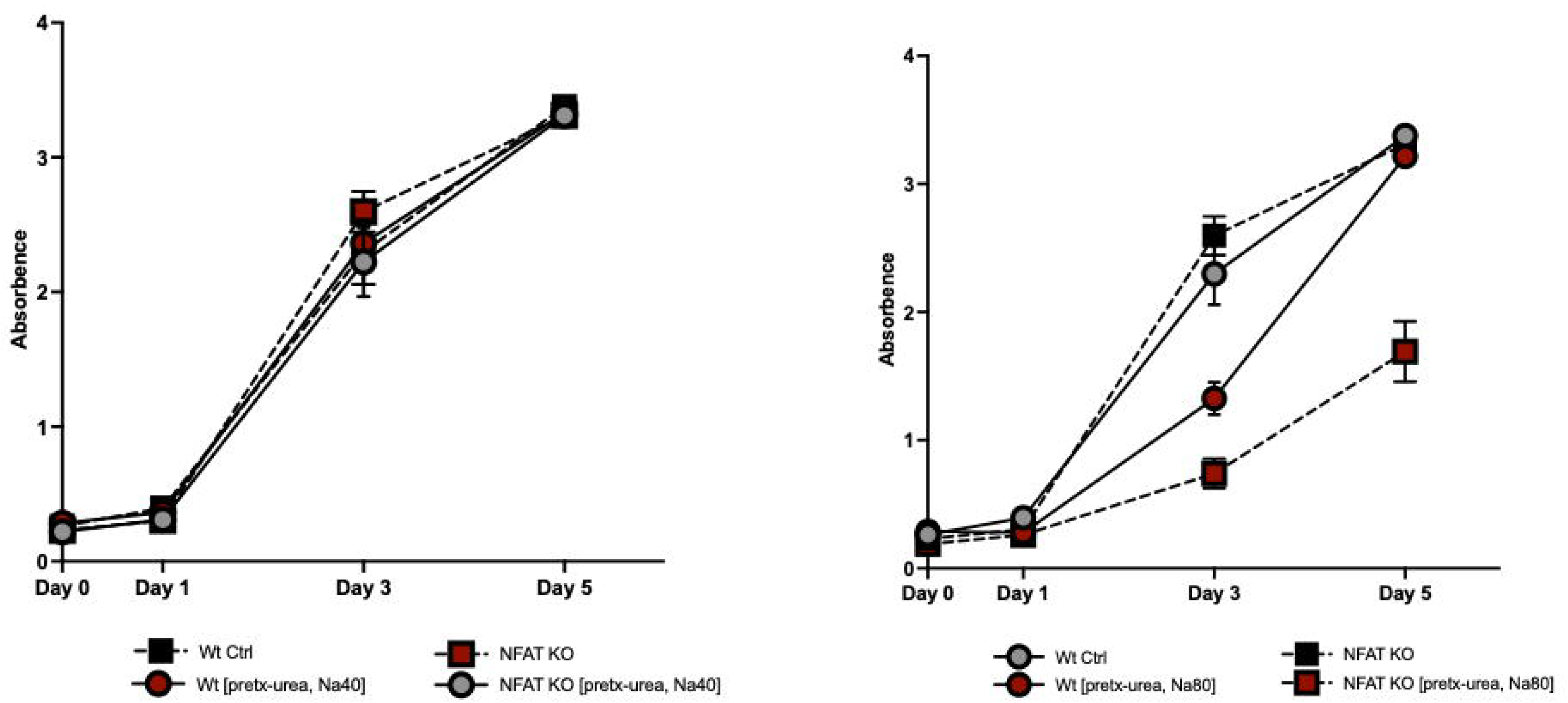
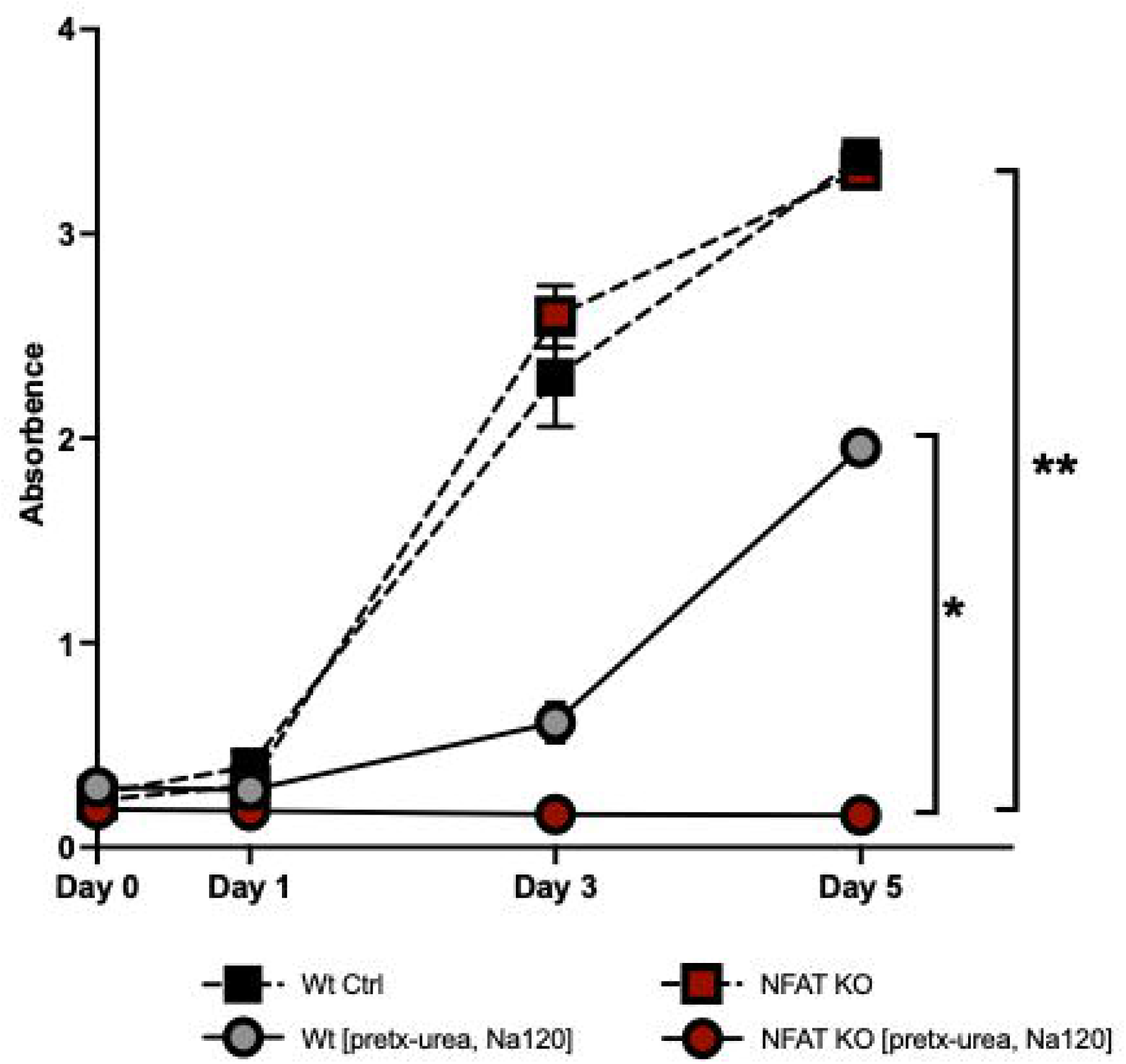
Urea priming induces alternate mechanisms for cell survival following hyperosmolar salt treatment. Cells were plated in media and grown in hyperosmolar urea (pretx-urea) using step-wise increases in osmolarity to a max of +80mM/80mOsms and allowed to acclimate. Cells were then treated with vehicle or additional osmoles of: (A) NaCl [+40mM/80mOsm], (B) NaCl [+80mM/160mOsm] or (C) NaCl [+120mM/240mOsm]]. Cell survival was determined at 0, 1, 3, or 5 days by staining with Janus Green B dye and measured at 620nm. spectrophotometer to determine concentration. Data were analyzed using a simple linear regression curve fitted to survival data followed by slope comparison, and are expressed as mean±SEM, n=4-5. *p<0.05, **p<0.01

### IMCD3 Renal Epithelial Cells Regulate Osmolyte Transporter Expression via NFAT5

To understand how IMCD3 cells respond to osmotic stress, we assessed mRNA expression in IMCD3 Wt and NFAT5 KO cells of genes known to be important in counteracting osmotic stress (Fig 4). We assessed expression of the betaine/GABA transporter (Fig 4A, *bgt1*) (14.36 ± 0.53 ΔCt Wt control *vs.* 22.05 ± 0.60 ΔCt NFAT5 KO control, ****p<0.0001, *2way* ANOVA, n=4), aquaporin 1 (Fig 4B, *aqp1*) (13.92 ± 0.28 ΔCt Wt control *vs.* 19.64 ± 0.60 ΔCt NFAT5 KO control, ****p<0.0001, *2way* ANOVA, n=4), and aldo-keto reductase 1B3 (Fig 4C, *akr1b3*) (6.17 ± 0.18 ΔCt Wt control *vs.* 7.72 ± 0.23 ΔCt NFAT5 KO control, ****p<0.0001, *2way* ANOVA, n=4). *Bgt1* and *aqp1* both encode for transport proteins which contribute to normalizing intracellular osmotic pressure by increasing amino acid and water transport, *resp. Akr1b3* encodes for aldose reductase, an enzyme which converts glucose to sorbitol and helps protect against osmotic pressure. Unsurprisingly, our data show considerably decreased baseline expression of these membrane transport proteins in IMCD3 NFAT KO cells, which has been shown for some in previous studies^5^. For both *bgt1* and *aqp1* there were trending increases in Wt expression with increasing osmolar stress. This hyperosmolar response was significant for *akr1b3* (6.17 ± 0.18 ΔCt Wt control *vs.* 4.24 ± 0.23 ΔCt Wt NaCl [80mM], ****p<0.0001 & 4.79 ± 0.23 ΔCt Wt control *vs.* 6.17 ± 0.18 ΔCt Wt mannitol [80mM], **p<0.01, *2way* ANOVA, n=4). However, IMCD3 NFAT5 KO cells were unable to similarly upregulate *akr1b3* in response to the osmotic stress.

**Figure 4:**
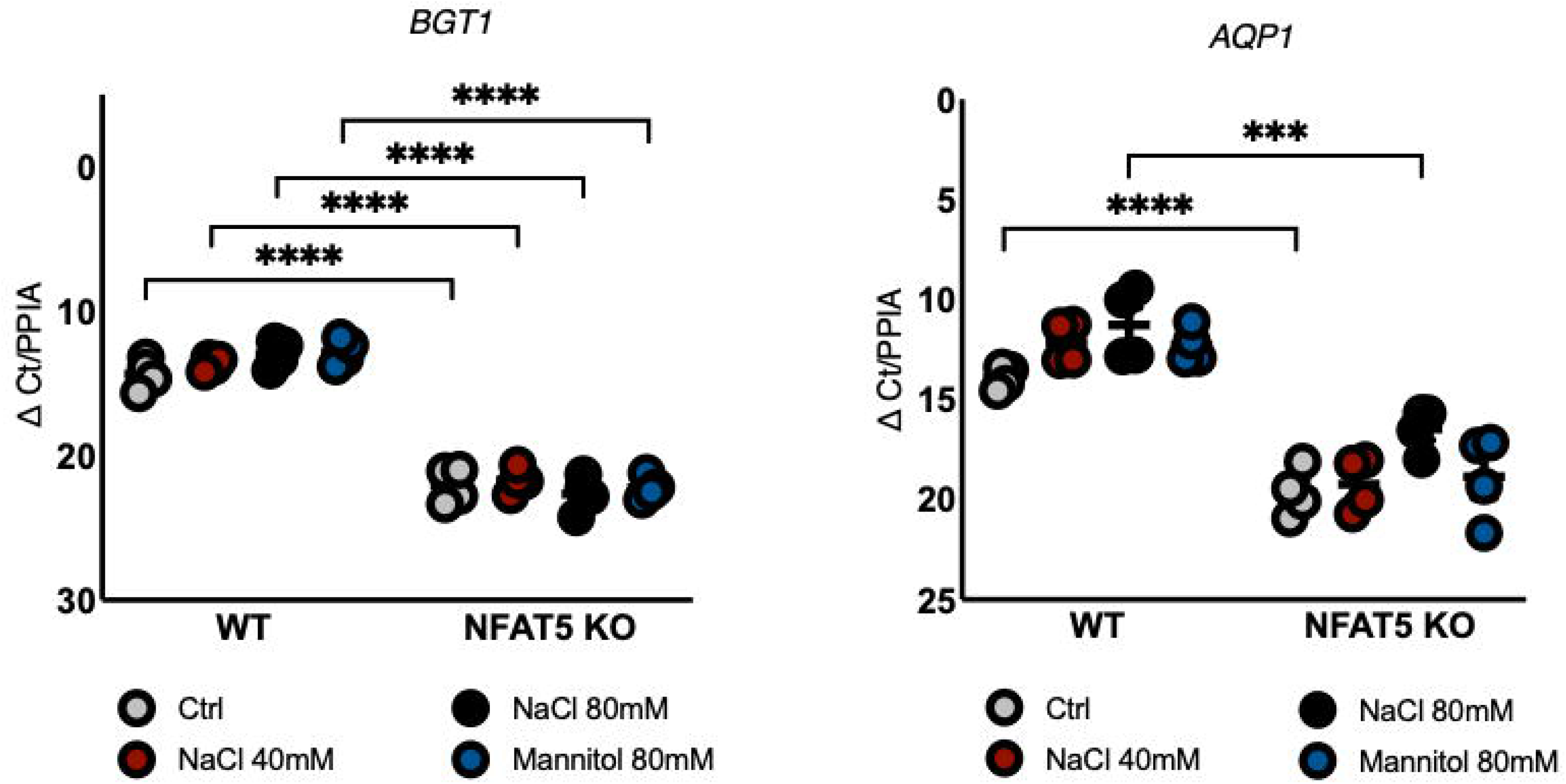
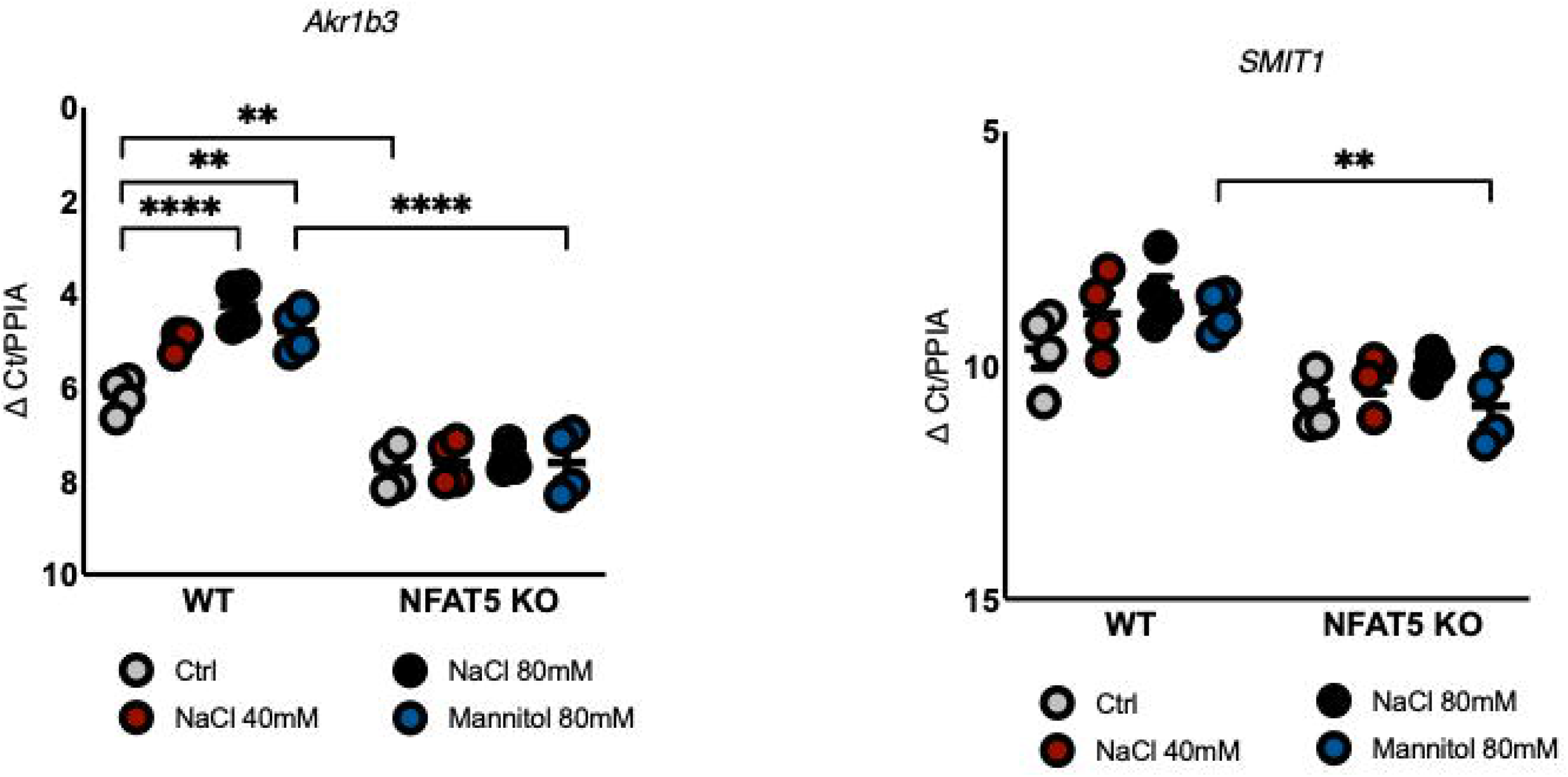
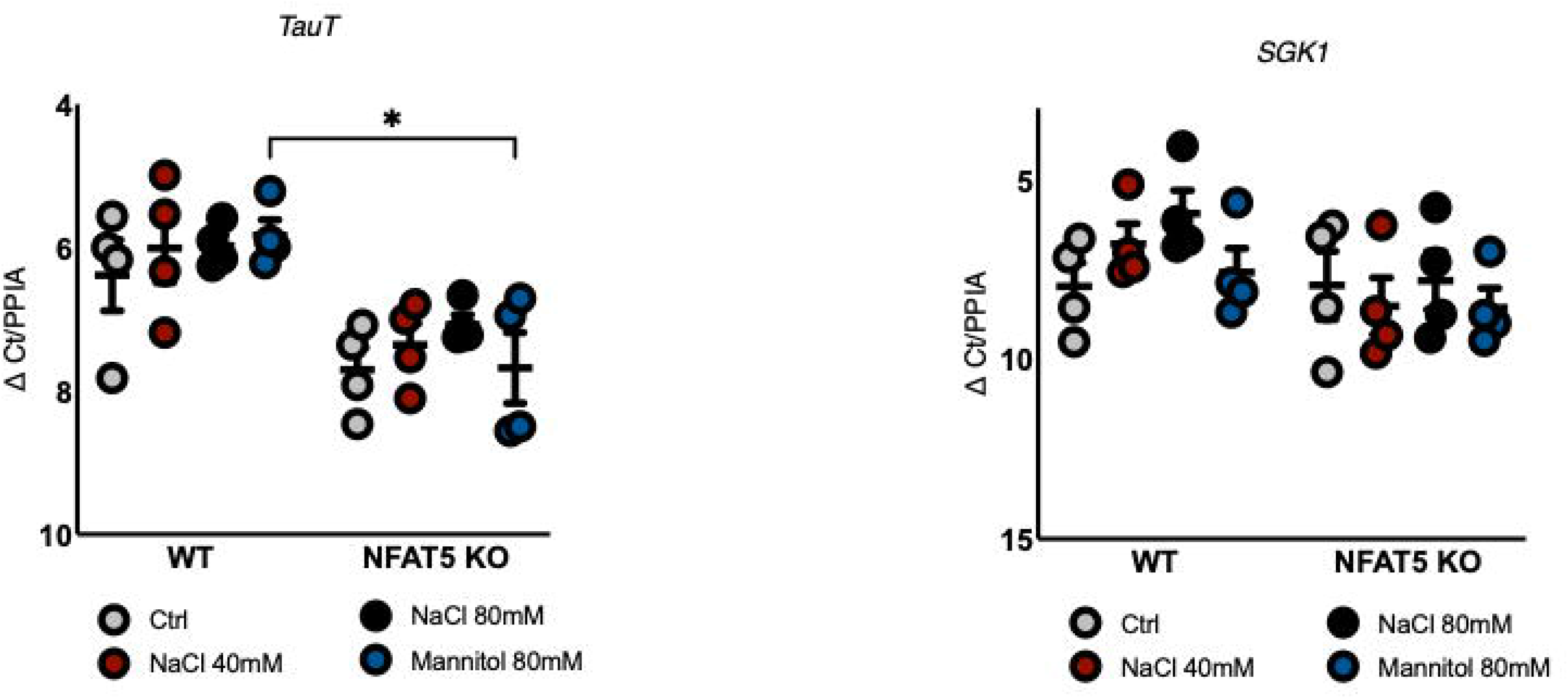
Nuclear factor of activated t-cells (NFAT5)-dependent gene expression in inner medullary collecting duct (IMCD3) cells. Wt or NFAT5 KO IMCD3 cells were treated with vehicle or additional osmoles of NaCl [+40mM/80mOsm], NaCl [+80mM/160mOsm] or mannitol [+80mM/160mOsm]. Total mRNA was isolated to determine gene expression of targets known to be affected with hyperosmolarity, including (A) *BGT1*, (B) *AQP1*, (C) *Akr1b3*, (D) *SMIT1*, (E) *TauT* and (F) *SGK1* expression. Cycle threshold (CT) values obtained and normalized to housekeeping (*PPIA*) gene expression. Data were analyzed using ΔCT/PPIA, and are expressed as mean±SEM, performed in triplicates (averaged), n=5. *p<0.05, **p<0.01, ***p<0.001, ****p<0.0001; ANOVA with Tukey’s posthoc

While not significant, sodium-myo-inositol transporter 1 (Fig 4D, *SMIT1)* and taurine transporter (Fig 4E, *TauT)*, expression showed a trending lower baseline expression in the NFAT5 KO cells compared to the Wt. Additionally, the Wt cells showed a trending increase for *SMIT1* expression in the treated groups as compared to control, while the NFAT5 KO cells did not.

Serum-glucocorticoid kinase 1 (*SGK1*) expression increases in response to increased osmolarity expression; however, we did not observe a lower baseline expression in the NFAT5 KO cells as compared to the Wt (Fig 4F). However, Wt cells did exhibit a trending increase in SGK1 expression with treatment, while NFAT5 KO cells did not. These differences may reveal why NFAT5 KO cells exhibited some ability to recover in the presence of osmotic stress. Together, these data confirm the critical role for NFAT5 in regulating osmolyte transporter and water channel expression in the face of osmotic stress (Fig 5) but also demonstrate possible redundant pathways that can prevent cell death.

**Figure 5:**
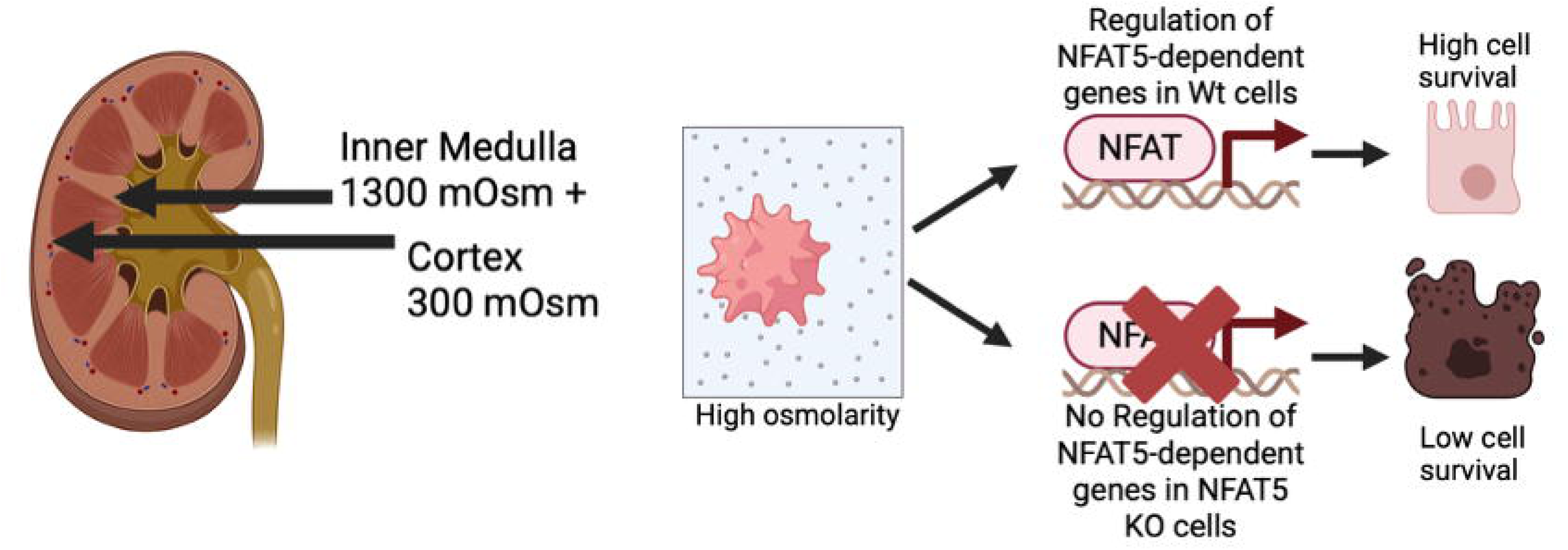
Nuclear factor of activated t-cells (NFAT5)-dependent increases in genes responsible for countering osmotic stress is crucial for cell survival in high osmolarity. Schematic showing that inner medullary collecting duct cells depend upon *NFAT5* expression to counter increasing hyperosmolar stress in the medulla. Cells without *NFAT5* exhibit reduced cell survival; however, urea priming does allow for better survival in hyperosmolar conditions.

Unfortunately, we were unable to develop a CRISPR NFAT5 KO RAW cell line using the same methods as our IMCD3 cell line. Instead, we used siRNA to knockdown NFAT5 (NFAT5 KD) expression (Fig 6A). Using lipofectamine, we observed a significant reduction in NFAT5 (7.544 ± 0.191 ΔCt NFAT5 KD *vs.* 6.536 ± 0.129 ΔCt scramble, ***p<0.001, *Students’ t-test*, n=12) but were never able to completely knock out NFAT5 expression. This limitation restricted our ability to perform chronic experiments in this model, as well as to completely compare with our IMCD3 cell line. Other papers show similar technical difficulties in modulating gene expression with siRNA in immune cell lines and observed similar levels of gene KD, without exhibiting cell toxicity^27,28^.

**Figure 6:**
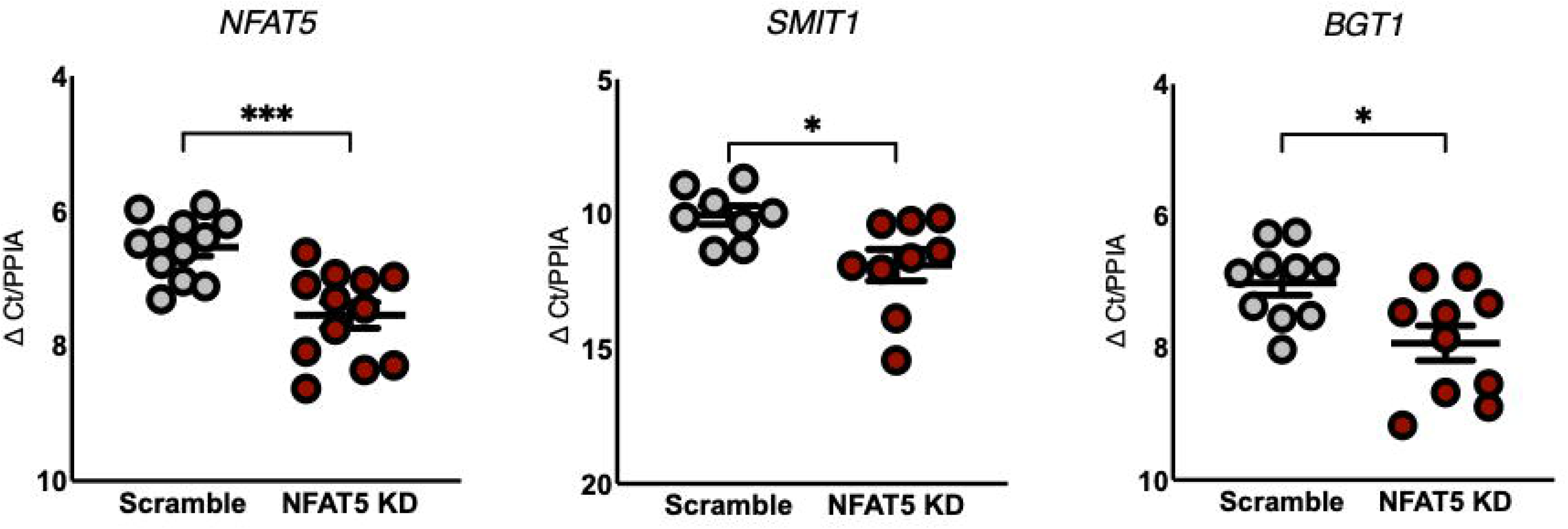
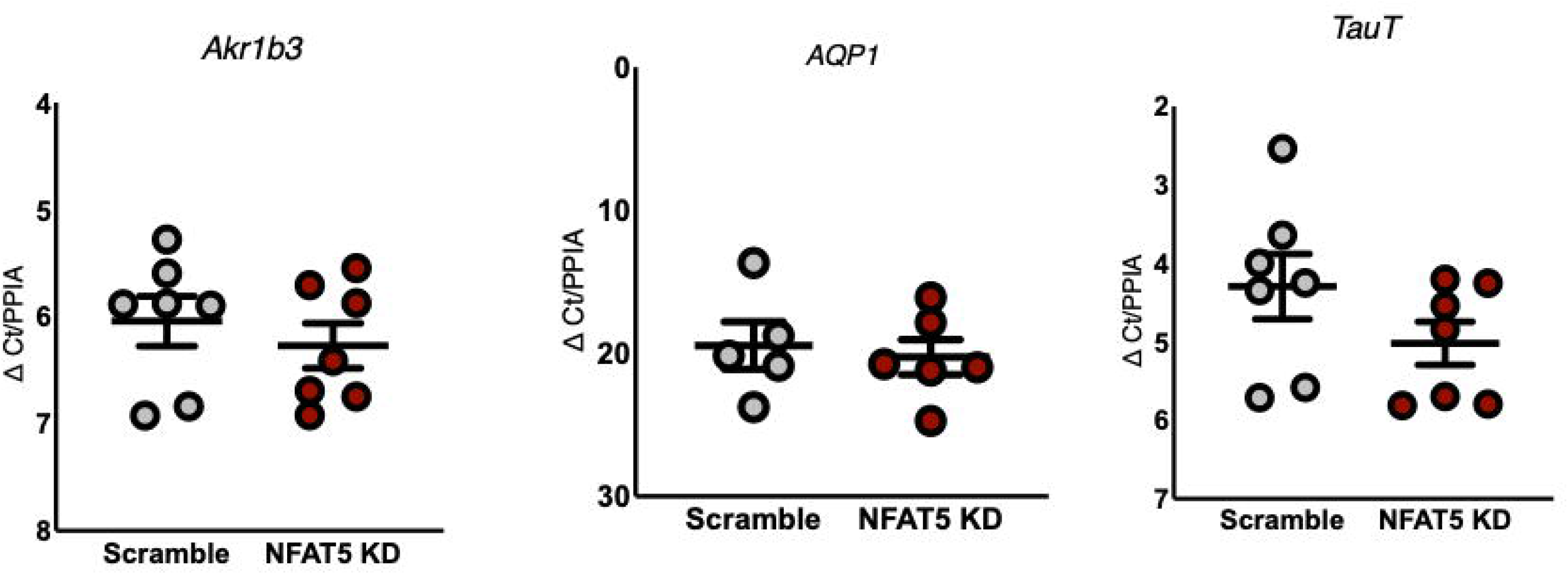
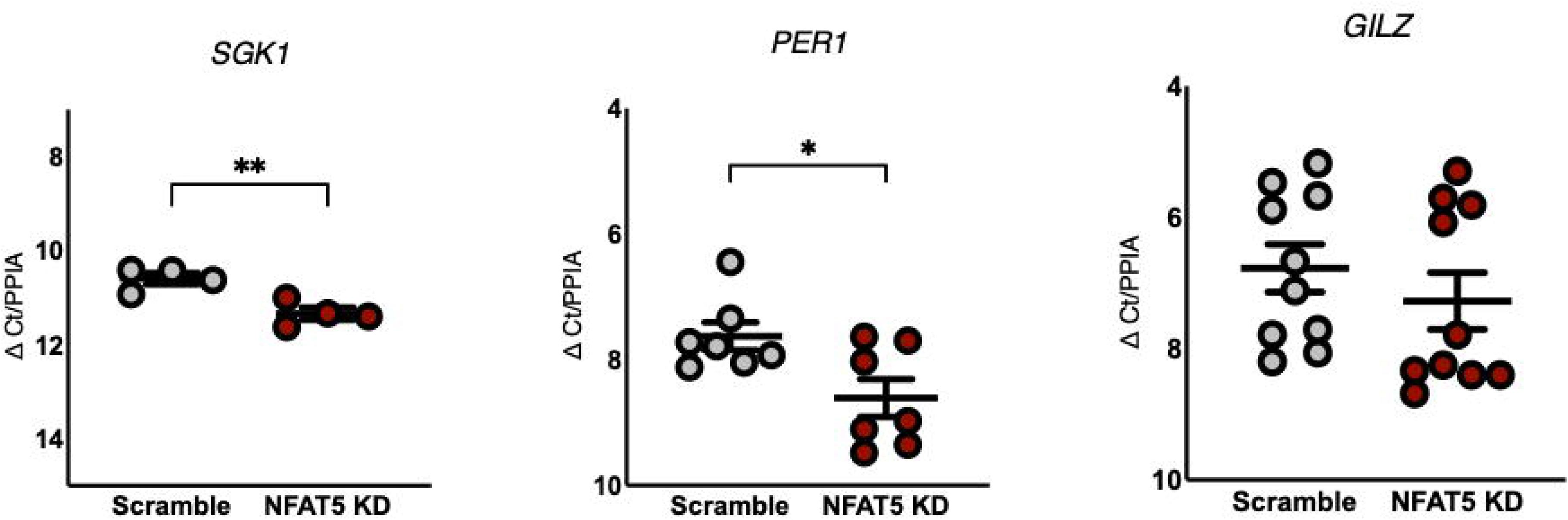
Nuclear factor of activated t-cells (NFAT5)-dependent gene expression of hyperosmolar stress response genes in RAW264 cells. RAW264 cells, a model of innate immune cells, were transfected with siRNA (A) against *NFAT5*. Cells showed partial reductions in *NFAT5* gene expression. Cycle threshold (CT) values obtained and normalized to housekeeping (PPIA) gene expression. Data were analyzed using ΔCT/PPIA, and are expressed as mean±SEM, performed in triplicates (averaged), n=12. ***p<0.001, *Students’ t-test*. Wt or NFAT5 knockdown (KD) RAW264 cells were grown in media and total mRNA was isolated to determine gene expression of targets known to be affected with hyperosmolarity, including (A) *SMIT1*, (B) *BGT*1, (C) *SGK1*, (D) *PER1* or (E) *GILZ*. Cycle threshold (CT) values obtained and normalized to housekeeping (PPIA) gene expression. Data were analyzed using ΔCT/PPIA, and are expressed as mean±SEM, performed in triplicates (averaged), n=4-9. *p<0.05, **p<0.01; *Students’ t-test*

To determine NFAT5-dependent responses in RAW cells at baseline, RAW cells were transfected with NFAT5 siRNA or scramble siRNA before extracting total RNA and performing qPCR analysis. RAW cells showed a significant decrease in gene expression for the two membrane transport proteins (Fig 6B, C): *SMIT1* (11.90 ± 0.59 ΔCt NFAT5 KD *vs.* 10.05 ± 0.35 ΔCt scramble, *p<0.05, *Students’ t-test*, n=8-9) and *BGT1* (7.93 ± 0.262 ΔCt NFAT5 KD *vs.* 7.02 ± 0.18 ΔCt scramble, *p<0.05, *Students’ t-test*, n=10). This mirrored results we observed in IMCD3 cells. Interestingly, in *Akr1b3* or *AQP1* levels were not reduced (Fig 6D, E) in RAW NFAT5 KD cells at baseline, which is different than responses observed in IMCD3 NFAT5 KO cells. *TauT*, which is a transporter for taurine, was also unchanged (Fig 6F). However, *SGK1* (11.34 ± 0.13 ΔCt NFAT5 KD *vs.* 10.60 ± 0.12 ΔCt scramble, **p<0.01, *Students’ t-test*, n=4) was decreased (Fig 6G) with NFAT5 KD. Given the role of SGK1 in circadian rhythm^29^, we also assessed NFAT5-dependent regulation of period circadian regulator 1 (*PER1*). We found that *PER1* was also decreased (Fig 6H, 8.61 ± 0.30 ΔCt NFAT5 KD *vs.* 7.63 ± 0.22 ΔCt scramble, *p<0.05, *Students’ t-test*, n=7). Yet, glucocorticoid-induced leucine zipper (*GILZ*), which responds to SGK-1, was not (Fig 6I) changed. These data highlight how NFAT5 may be differentially regulating proteins following osmotic stress.

### RAW Cells Regulate Cytokine Expression via NFAT5

Others have observed hyperosmolarity induced pro-inflammatory responses in immune cells, including macrophages^21,30^. To determine whether NFAT5 may be regulating pro-inflammatory cytokine expression, we assessed gene expression of interleukin 6 (*IL6*), tumor necrosis factor α (*TNFα*) and interleukin 1 beta (*IL1Δ*) following NFAT5 KD. We observed a significant reduction at baseline in both *IL6* (20.01 ± 0.26 ΔCt NFAT5 KD *vs.* 18.09 ± 0.18 ΔCt scramble, ***p<0.001, *Students’ t-test*, n=4-6) and *TNFα* (7.81 ± 0.09 ΔCt NFAT5 KD *vs.* 7.34 ± 0.14 ΔCt scramble, *p<0.1, *Students’ t-test*, n=9-10), but no differences in *IL1Δ* (Fig 7A, B, C). Together, these data suggest that NFAT5 may differentially regulate membrane transporters and proteins important for osmotic stress as well as cytokines in immune cells *vs.* epithelial cells (Fig 8).

**Figure 7:**
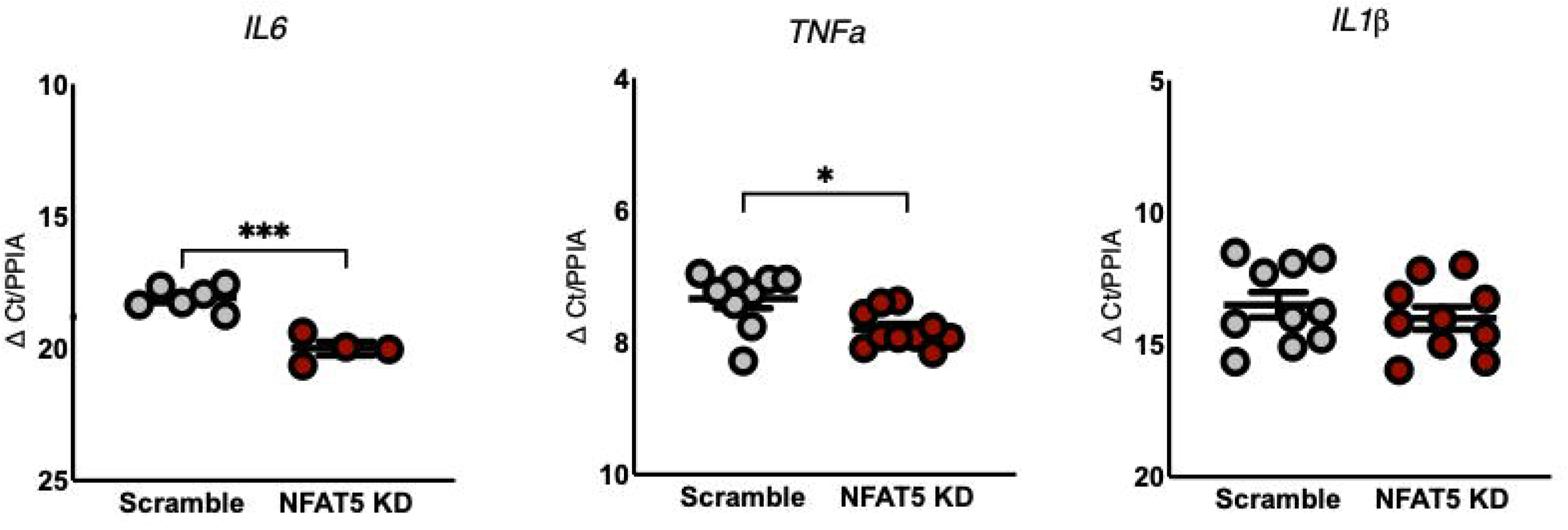
Nuclear factor of activated t-cells (NFAT5)-dependent cytokine gene expression RAW264 cells. Wt or NFAT5 knockdown (KD) RAW264 cells were grown in media and total mRNA was isolated to determine gene expression of cytokines produced by innate immune cells (A) *IL6*, (B) *TNFα*, or (C) *IL1Δ*. Cycle threshold (CT) values obtained and normalized to housekeeping (PPIA) gene expression. Data were analyzed using ΔCT/PPIA, and are expressed as mean±SEM, performed in triplicates (averaged), n=4-10. *p<0.05, ***p<0.001; *Students’ t-test*

**Figure 8:**
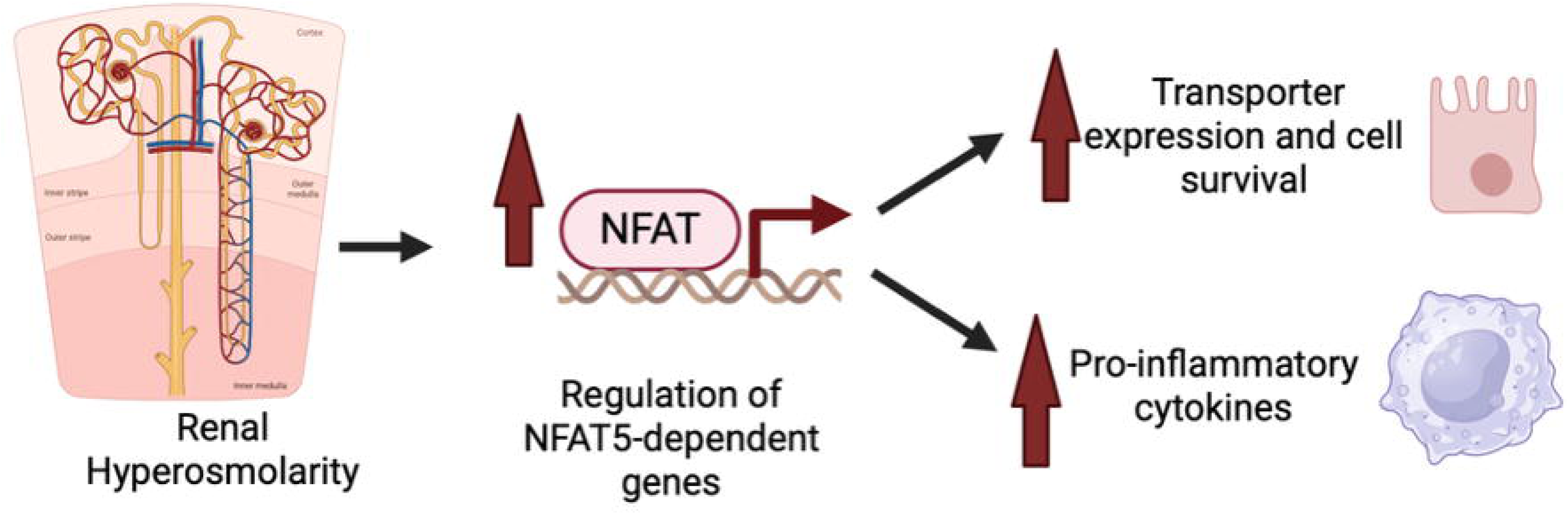
Differential gene regulation by nuclear factor of activated t-cells (NFAT5) Schematic showing that two predominate cells of the medulla-inner medullary collecting duct cells and immune cells- show differential responses to *NFAT5* gene expression.

## Discussion

Other investigators have used whole animal and cell-specific NFAT5 KO animal models^5,15,31^, but we chose to use *in vitro* models to specifically assess NFAT5-mediated regulation in different cell types found in the kidney which would be exposed to hyperosmolarity. Previous studies investigating the role of NFAT5 have shown crucial regulation of membrane transporters important for intracellular accumulation of osmolytes in the presence of extracellular hyperosmolarity^32^.

Here, we found that renal epithelial cells of the IM (IMCD3) are capable of survival in hyperosmolar media and that NFAT5 is critical for survival in this hostile environment. This study also suggests multiple NFAT5-dependent cell survival mechanisms which are not only reduced at baseline but also exhibit reductions with hyperosmolarity. Although not explored here, we found that renal epithelial cells diverted to another, albeit slower, mechanism for cell survival which does not rely on NFAT5. To propose targets responsible for the difference in survival between Wt and NFAT5 KO epithelial cells, we performed qPCR and showed that key transporters are upregulated when stimulated with hyperosmolarity predominantly in the Wt cells. However, NFAT5 KO cells did show a less pronounced change in expression of some key transporters when stimulated with hyperosmolarity, again suggesting a second mechanism that responds to hyperosmolarity without NFAT5.

We observed enhanced survival when the cells were grown in a hyperosmotic environment of urea, which has been explored before but with conflicting data. Lee and colleagues used a short, moderate urea treatment and found Madin-Darby canine kidney (MDCK) cells resistant to hyperosmolarity in subsequent treatments^33^. This corresponds with other reports showing that progressive increases in hyperosmolarity boosts survival, yet this response was only seen with hyperosmolar salt priming *vs.* hyperosmolar urea^34,35^. Here, we specifically investigated the role of NFAT5 in mediating urea priming-induced survival. We found that high NaCl [+80mM] significantly reduced cell proliferation in NFAT5 KO cells, but the priming before this stressful hypertonic stimulus allowed the cells to compensate. This reveals the existence of powerful NFAT5-independent osmo-adaptive mechanisms. There are multiple compensatory pathways, including general control nonderepressible (GCN)2 and activating transcription factor 4 (ATF4)^36–38^. For example, ATF4 drives expression of amino acid transporters^37,38^. This metabolic reprogramming may enhance cellular stress resilience through mechanisms completely independent of NFAT5-mediated osmolyte transporter induction and may be a cause of renal epithelial cell survival with urea priming.

Here, we also showed that NFAT5 has a divergent mechanism in epithelial *vs.* immune cells. In epithelial cells, NFAT5 acts to upregulate key transporters, while in immune cells NFAT5 promotes the expression of pro-inflammatory cytokines. This suggests NFAT5 could have additional divergent activity in other cell types exposed to hyperosmolarity. The observation that NFAT5 protects epithelial cells while promoting inflammation in macrophages reflects fundamental differences in cellular stress responses and transcriptional programs between these cell types^16,17^. Recent studies reveal that reactive oxygen species (ROS) is critical in directing which gene response NFAT5 directs. In addition, the source of ROS can specifically determine NFAT5 transcriptional specificity^20^. In renal epithelial cells under hypertonic stress, mitochondrial-dependent ROS production promotes NFAT5 nuclear localization, directing it toward osmo-protective gene promoters including heat-shock protein 70 (HSP70), osmolyte transporters, and aquaporins^20,39^. This mitochondrial ROS signature enables survival in the hyperosmotic renal medulla. In contrast, macrophages activated by toll-like receptor (TLR) ligands generate ROS through xanthine oxidase activating MAPK, recruiting NFAT5 to inflammatory gene promoters such as: IL-6, TNF-α and inducible nitric oxide synthase (iNOS)^20,21^. Beyond ROS sources, cell-type specific co-factor interactions shape NFAT5 function. In macrophages, NFAT5 and nuclear factor kappa-light-chain-enhancer of activated B cells (NF-κB) form synergistic partnerships^16,40^. In epithelial cells, NFAT5 functions more autonomously through direct binding to tonicity-responsive elements as a homodimer^3,18^. Our findings show differential expression of *BGT1*, *AQP1*, and *Akr1b3* between IMCD3 and RAW cells, with these genes highly NFAT5-dependent in epithelial cells but not in macrophages, directly support this mechanistic framework. The observation that RAW NFAT5 knockdown reduced *SMIT1* and *BGT1* but not *Akr1b3* or *AQP1* suggests that immune cells utilize only a subset of osmo-protective pathways, potentially reflecting their evolutionary adaptation to transiently patrol hypertonic tissue environments rather than permanently reside there^16,41^.

NFAT5-deficient mice exhibit severe T cell lymphopenia, suggesting a role for immune cell homeostasis and lymphoid organ development^42^. We and others show that NFAT5 can be pro-inflammatory in immune cells, with further studies showing increased cytokine secretion with hyperosmolar sodium^12,16,20^. Under high NaCl conditions, T cell differentiation skews toward a pathogenic Th17 phenotype through a NFAT5-SGK1 pathway^12,42^. Recent studies suggest that there can be an accumulation of tissue sodium, although whether that occurs with edema is not known^43–48^. However, there is a correlation with this increase in sodium and inflammation, as well as subsequent development of cardiovascular and organ damage^49–51^. These data suggest that increased sodium, via NFAT5, may be able to activate immune cells and increase inflammation, demonstrating a context specific role for NFAT5.

These mechanistic findings provide a framework for interpreting the differential responses of IMCD3 epithelial cells *vs* RAW cells to reductions in NFAT5 expression. The epithelial cells require NFAT5 for osmolyte transporter induction and cannot survive high NaCl without this transcriptional program. However, urea priming seems to create a pre-adapted-state that enables partial tolerance to hyperosmolar stress. This may also explain the physiological wisdom of using both NaCl and urea in the renal medulla; they activate complementary protective mechanisms providing redundancy against cell death^33,52^. However, macrophages show fundamentally different NFAT5 biology because they operate in different microenvironments with different primary stressors which are directed towards inflammatory rather than osmo-protective function.

## Acknowledgements

The authors would like to thank Masaaki Yoshigi, MD for technical assistance.

